# Chemodiversity in flowers of *Tanacetum vulgare* has consequences on a florivorous beetle

**DOI:** 10.1101/2023.06.19.545548

**Authors:** Rohit Sasidharan, Lukas Brokate, Elisabeth J. Eilers, Caroline Müller

## Abstract

The chemical composition of plant individuals can vary, leading to high intraspecific chemodiversity. Diversity of floral chemistry may impact the responses of flower-feeding insect visitors.
Plants of *Tanacetum vulgare* vary significantly in their leaf terpenoid composition, resulting in distinct chemotypes. We investigated the composition of terpenoids and nutritents of flower heads and pollen in plants belonging to three chemotypes, dominated either by β-thujone (BThu), artemisia ketone (Keto) or a mixture of (*Z*)-myroxide, santolina triene and artemisyl acetate (Myrox). Moreover, we tested the effects of these differences on preferences, weight gain and performance of adults of the shining flower beetle, *Olibrus aeneus*.
The terpenoid composition and diversity of flower heads and pollen significantly differed among individuals belonging to these chemotypes, while total concentrations of terpenoids, sugars, amino acids or lipids of the pollen did not differ. Beetles preferred the BThu over the Myrox chemotype in both olfactory and contact choice assays, while the Keto chemotype was marginally repellent in olfactory assays. The beetles gained the least weight within 48 h and their initial mortality was highest when feeding exclusively on floral tissues of the Myrox chemotype. Short-term weight gain and long-term performance were highest on the BThu chemotype.
In conclusion, the beetles showed chemotype-specific responses towards different *T. vulgare* chemotypes, which may be attributed to the terpenoid composition in flower heads and pollen rather than to differences in nutrient profiles. Both richness and overall diversity are important factors when determining chemodiversity of individual plants and their consequences on interacting insects.

**SHORT SUMMARY:** We demonstrate that *Tanacetum vulgare* chemotypes distinguished by their leaf terpenoid profiles also show unique floral and pollen chemotypes based on terpenoid composition and diversity, which affect the preference and performance of a beetle florivore.

## INTRODUCTION

Plant chemodiversity refers to the chemical diversity of specialized (secondary) metabolites, encompassing their richness, relative abundance and eveness (Müller and Junker, 2022). High chemodiversity is not limited to inter-species variation, but can also occur within species. Such intraspecific chemodiversity plays an important role in various biotic and abiotic interactions (Petrén et al., 2023; Wetzel and Whitehead, 2020). In several plant species, diversity is particularly found in distinct classes of specialized metabolites, such as glucosinolates, glycoalkaloids or terpenoids (Calf et al., 2019; Kleine and Müller, 2011; Tewes and Müller, 2020). Based on these classes, different chemotypes can be distinguished, which are dominated by one or several of these metabolites. Previous research has mainly focused on investigating differences in the chemical profiles of these chemotypes in relation to leaves, studying their consequences on herbivorous invertebrates and pathogens (Salazar et al., 2016; Tewes and Müller, 2020; Whitehead and Bowers, 2013; Ziaja and Müller, 2023). In contrast, limited knowledge exists regarding chemotypic variation in flowers and its consequences for flower-visiting insects (Egan et al., 2018; Eilers et al., 2021; Kessler et al., 2013; Rusman et al., 2022).

Flowering plants interact with a large range of animals, including pollinators, herbivores or predators. The flowers produce numerous specialized metabolites, which serve various functions in these interactions. Ideally, volatile plant metabolites should attract pollinators from a distance, while repelling antagonists, such as herbivores (Strauss and Whittall, 2006; Theis and Lerdau, 2003). However, floral herbivores, known as florivores, also exploit these volatiles for host plant finding (Junker and Blüthgen, 2010; Knauer and Schiestl, 2015; Schiestl, 2010; Sasidharan et al., 2023). Terpenoids represent a major class of specialized metabolites that are very diverse in structure and occur in different plant parts, including the flowers (Abbas et al., 2017; Zhou and Pichersky, 2020). Many terpenoids are volatile and contribute thus to the floral scent. Plant individuals of the same species that differ in their floral volatile profiles, including the terpenoids, may result in different flower visitation behaviour. The chemical composition of the floral blend is thereby highly decisive for the first host plant choice of insect visitors (Junker and Parachnowitsch, 2015; Schiestl, 2010).

Once landed on the flowers, the nutritional quality, determined by the presence of nutrients as well as specialized metabolites in the floral parts, influences the feeding behaviour and in the long-term the developmental performance and survival of flower-visitors. Floral parts such as pollen, which serve as rewards for pollinators, are rich in amino acids, sugars, lipids and several micronutrients (Vaudo et al., 2016). For example, the amino acids L-alanine and L-serine induce electroantennographic responses and preference behaviour in the western corn rootworm, *Diabrotica virgifera virgifera* (Coleoptera: Chrysomelidae) (Hollister and Mullin, 1998). Likewise, different sugars, such as fructose, glucose, sucrose and isorhamnetin rhamnoglycoside, act as phagostimulants for this florivore (Kim and Mullin, 2007). However, plants also incorporate deterrent or toxic compounds, including terpenoids, in these tissues to defend against overexploitation by pollinators or florivores (Palmer-Young et al., 2019; Rivest and Forrest, 2020; Sasidharan et al., 2023). For example, a glucosinolate and saponins act deterrent against the pollen beetle, *Brassicogethes aeneus* (Coleoptera: Nitidulidae), in oilseed rape (Austel et al., 2021). The nutrient content in pollen may be correlated to its specialized metabolite diversity (Sasidharan et al., 2023), but differences in nutrient composition among plants of different chemotypes have rarely been investigated (Eilers et al., 2021). The selection of a suitable host plant is also critical for the survival of florivores. Adult pollen beetles showed distinct survival times when fed with pollen of different genotypes of oilseed rape differing in host plant quality (Hervé et al., 2016).

*Tanacetum vulgare* (Asteraceae), common tansy, is a highly chemodiverse species with a large geographic distribution that is attacked by various herbivorous species (Kleine and Müller, 2011; Schmitz, 1998; Wolf et al., 2012). It shows a particularly high terpenoid richness and concentration in the leaves and flower heads, based on which individual plants can be assigned to different chemotypes (Keskitalo et al., 2001; Kleine and Müller, 2011; Rohloff et al., 2004). These chemotypes can be either mono chemotypes, whose terpenoid profile is dominated (>40%) by one terpenoid or mixed chemotypes with two or more terpenoids being predominant (Holopainen et al., 1987). The chemotype identity of *T. vulgare* has been reported to influence the distribution and behaviour of various leaf herbivores, particularly aphids (Benedek et al., 2015; Clancy et al., 2016; Jakobs et al., 2019; Ziaja and Müller, 2023) but also beetles, lepidopterans and leaf miners (Kleine and Müller, 2011; Susurluk et al., 2007; Wolf et al., 2012). Plants of different (leaf) chemotypes were also found to differ in various floral traits, including the protein content and the protein to lipid ratio in pollen (Eilers et al., 2021). In a semi-field experiment with plots consisting of either one or different mono-chemotypes of *T. vulgare*, less individuals of the shining flower beetle, *Olibrus* sp. (Coleoptera: Phalacridae), were found in the latter, indicating that the overall diversity is important for their host choice (Eilers et al., 2021). Adults of *Olibrus* species mainly feed on pollen, but also oviposit in the flower heads of Asteraceae (Gaafar et al., 2016). However, it was yet unknown whether intraspecific plant chemodiversity influences the beetles’ preference and performance.

Thus, we investigated whether the flowers of plants of three chemotypes, defined by their leaf terpenoid profiles, also differ in their terpenoid and pollen nutrient composition, i.e., sugars, amino acids and total lipids. We used two mono-chemotypes, dominated either by artemisia ketone (henceforth Keto) or β-thujone (BThu), as well as a mixed chemotype, containing (*Z*)-myroxide, santolina triene and artemisyl acetate (Myrox), showing increasing levels of Shannon diversity of their leaf terpenoids (Ziaja and Müller, 2023) and described their diversity with Renyi profiles (Tóthmérész, 1995). Moreover, we studied potential consequences on the preference behaviour of beetles of *Olibrus aeneus* and their performance in the laboratory. We hypothesized that (1) *T. vulgare* chemotypes differ in their floral terpenoid and nutrient composition, with terpenoid diversity in both flower heads and pollen reflecting the leaf terpenoid diversity and terpenoid-rich pollen containing more nutrients. Furthermore, we hypothesized that (2) beetles have a lower preference for the more diverse chemotype, and (3) beetle performance is lower on both low-(due to poorer nutrients) and high-(due to toxin effects) diversity chemotypes than on the intermediate-diversity chemotype, which should have a balanced nutrient content and diversity of terpenoids.

## METHODS

### Plant and insect material

A stock of *T. vulgare* plants was established in the greenhouse from seeds that had been collected in Bielefeld, Germany. Chemotypes had been determined based on the leaf terpenoid profiles (for further details, see Eilers et al., 2021; Ziaja and Müller, 2023). Plants were propagated through cuttings to produce clonal lineages that maintain their respective chemotypes. The cuttings were made from plant individuals (called maternal origins here) belonging to the three chemotypes, dominated either by artemisia ketone (Keto, average Shannon index (H_s_) of leaf terpenoids = 1.15, seven maternal origins in total), β-thujone (BThu, average H_s_ = 1.26, eight maternal origins) or a mixture of (*Z*)-myroxide, santolina triene and artemisyl acetate (Santo, average H_s_ = 2.16, six maternal origins). Plants were maintained in 4 L pots in a greenhouse at a 16 h: 8 h light: dark cycle and temperatures around 20°-22 °C and fertilized weekly (modified solution according to Arnon and Hoagland, 1940). For chemical analyses, freshly blooming flower heads were collected, flash-frozen in liquid nitrogen and stored at −80 °C until further processing. Pollen was collected by shaking freshly open flower heads on wax-coated paper, from which the pollen was transferred into Eppendorf tubes on ice and stored immediately at −20 °C for bioassays and chemical analyses. For the latter, pooled pollen samples from the same plant individual were divided in subsamples for different analyses.

Beetles of *O. aeneus* were collected in Bielefeld, Germany (52°02’48.4’’ N, 8°29’30.4’’ E), from flowers of *Matricaria reticulata* and other Asteraceae plants between July and September 2021 (for beetle identification see Supplement Fig. S1). They were maintained in plastic boxes (22 cm x 13 cm x 12 cm) with freshly provided flower heads from *T. vulgare* belonging to a mixture of the different chemotypes for at least 1-2 weeks before they were used in the bioassays. These boxes were kept in a climate cabinet at 22 °C and 60% relative humidity, under a 16:8 light:dark cycle. Oviposition was not observed in the laboratory and thus only field-collected adult beetles were used in all of the bioassays within eight weeks after collection.

### Chemical analyses of terpenoids from flower heads and pollen, sugars and amino acids from pollen

Terpenoids of flower heads and pollen were conducted similarly as in Eilers et al. (2021). Flower heads were freeze-dried and homogenized. Around 10 mg of the sample was extracted with 1 mL of *n*-heptane containing 10 ng μL^-1^ 1-bromodecane (Sigma-Aldrich, Germany) as internal standard, homogenized, ultra-sonicated for 5 min and then centrifuged. Supernatants and blanks containing the internal standard only were analyzed via gas chromatography coupled to a mass spectrometer (GC-MS, GC-2010 Plus coupled to QP2020; Shimadzu, Kyoto, Japan) with a VF-5 MS column (Agilent Technologies, Santa Clara, CA, USA; 30 m x 0.25 mm x 0.25 µm, with circa 10 m guard column). One μL of the sample was injected in a 10:1 split mode at 240 °C with a carrier gas (He) flow of 1.14 mL min^-1^. The oven was held at 50 °C for 5 min, increased to 250 °C at 10 °C per min and finally to 300 °C at 30 °C per min, then held at that temperature for 10 min. Compounds were identified through comparison of the retention index (RI) and mass spectra with library entries of the National Institute of Standards and Technology NIST 2014, Pherobase (El-Sayed, 2012) and mass spectra reported in Adams (2007). Terpenoid peaks were quantified by dividing the extracted ion peak areas by the area of the internal standard and the sample weights (dry weights for flower heads and fresh weights for pollen).

For sugar analyses of pollen, 10 mg of each sample was extracted in a mixture of 5:2:2 methanol:deionized water:chloroform containing ribitol (99%, AppliChem, Darmstadt, Germany) as internal standard, homogenized, 380 µL of deionized water was added and the sample was vortexed and centrifuged. The supernatants were derivatized with *O*-methylhydroxylamine hydrochloride (>97%; Sigma Aldrich-Merck, Madrid, Spain; >98%; Alfa Aesar, Kandel, Germany) in pyridine (≥99.9%, HPLC grade; Sigma-Aldrich, Germany) for 30 min at 50 °C. Afterwards, samples were silylated with N,O-bistrifluoroacetamide (Sigma-Aldrich, Germany) at 50 °C for 30 min. Samples were diluted in pyridine and analyzed by GC-MS. Blank runs and controls containing only the reaction mixture were also analyzed. Samples were injected at 240 °C, the oven was held at 100 °C for 1 min, increased to 200 °C at 2 °C per min, followed by an increase to 280 °C at 10 °C per min and a 5 min hold, and finally to 300 °C at 20 °C per min, then held at that temperature for 5 min. Analytes were identified based on their RI, mass spectra and mass-to-charge ratios (*m*/*z*) by comparing these with entries in the GOLM Metabolome database (Hummel et al., 2010; Kopka et al., 2005), mass spectral and retention time index libraries (Schauer et al., 2005) and with commercially available standards measured with the same method. The analytes were quantified based on their peak heights (total ion counts) and analyte peaks of fructose and glucose summed up, which each had two peaks. Peak intensities were related to the internal standard and the sample fresh weight.

For amino acid analyses of pollen, 10 mg of the pollen sample was digested with 6N HCl (Fisher-Scientific, UK) and boiled at 100 °C for 4 h to release bound amino acids. After cooling, samples were centrifuged and supernatants dried. These samples were extracted twice in 80% MeOH, containing norvaline and sarcosine (Sigma-Aldrich, Germany) as internal standards and analyzed after derivatization as in Jakobs and Müller (2018) by a high performance liquid chromatograph coupled to a fluorescence detector (HPLC-FLD; 1260/1290 Infinity HPLC und FLD; Agilent Technologies) equipped with a ZORBAX Eclipse Plus C18 column (250 x 4.6 mm, 5 μm particle size, with guard column; Agilent Technologies). The chromatographic separation was performed using a gradient from eluent A to eluent B, starting at 2% B [4.5 acetonitrile (LC-MS grade; Thermo Fisher Scientific): 4.5 methanol: 1 Millipore water, *v*:*v*:*v*], held for 0.84 min and then increased linearly to 57% B (reached at 68.4 min), followed by a cleaning and equilibration cycle. The flow rate was 1.5 mL min^-1^ and the column temperature 40 °C. Amino acid derivatives were detected with excitation wavelengths of 340 nm for primary and 260 nm for secondary amino acids and emission wavelengths of 450 nm and 325 nm, respectively. Blank runs were also performed. Data were analyzed using OpenLab ChemStation C.01.07 (Agilent Technologies). Amino acids were identified by RT matching using the standard mixes and quantified by their peak heights in relation to the internal standards (norvaline for primary, sarcosine for secondary amino acids). Peak intensities were related to sample fresh weight.

Lipids were extracted from 2 mg dried pollen samples by homogenization in 200 μL 2% Na_2_SO_4_. One mL of chloroform:methanol (1:1 v:v) was added, samples sonicated for 15 min, and water added for phase separation. A concentration series was prepared in chloroform from dodecanoic acid (Sigma-Aldrich, Merck, Germany). Samples were dried under vacuum at 60 °C and resolved in 200 μL 98% sulfuric acid. Of each sample and standard, 50 μL triplicates were applied on microtiter plates, sealed and incubated for 10 min at 95 °C. After incubation for 5 min at 4 °C, plates were unsealed and 100 μL of 400 μg mL^-1^ vanillin in 34% phosphoric acid was added to each well. After 10 min at room temperature, the absorbance was measured at 570 nm. Total lipid concentrations were calculated based on the dodecanoid concentration series, averaging the three replicate values per sample.

### Olfactometer choice assay

A glass Y-tube olfactometer was used to study the olfactory preference of *O. aeneus* beetles between different sources (Fig. S2). The Y-tube had two side arms of 10 cm in an angle of 45° and a central arm of 15 cm, with a diameter of 2 cm. Air was pumped at a continuous flow rate of 300-400 mL min^-1^ through the two side arms. The Y-tube was placed in a cardboard box covered with white paper to prevent visual cues, and angled at a slope of 45 ° with the help of a ramp also covered with white paper. A fluorescent lamp (∼80 μE m^-2^) was placed above the setup to stimulate beetle activity. Glass cylinders were connected to the two side arms to provide odour cues. Two to three flower heads were placed inside these glass cylinders as odour sources, depending on the size of the flower heads. Beetles were offered the choice between a) flower heads of one of the chemotypes against a blank odour control and b) flower heads of two different chemotypes (in total three combinations).

Beetles were prevented from leaving the Y-tube by a Teflon mesh covering both ends of the side arms. Beetles were starved for 16-24 h prior to the assay. The position of the Y-tube was switched after every few runs to minimize bias in the observations and the Y-tube was cleaned with ethanol after every four runs and dried in an oven. The flower heads were also replaced after every four runs. Beetles were allowed to make a choice within five minutes. If they did not make a choice within that time they were considered as non-responders. A response was counted when a beetle entered one of the side arms of the tube. In total, at least 34 responsive beetles were tested per combination. Each beetle was tested only once per combination.

### Contact choice assay

To test the preference of beetles for certain chemotypes in a contact assay, we used a small tube made of polytetrafluoroethylene (4 cm long, 0.5 cm diameter), illuminated from above with a fluorescent lamp (∼80 μE m^-2^). The set-up was filmed with a camera placed about 10 cm above the tube. Beetles were entered through a hole (o.5 cm diameter) in the center of the tube. One flower head of two chemotypes was placed at the end of either side of the tube (three combinations of the chemotypes in total). Beetles were starved for 24 h prior to the assay. After every trial, the tube with flower heads was rotated by 90° and flower heads were replaced every two trials. When a beetle contacted a flower head for at least 10 s, it was considered to have made a contact choice. In addition, the contact duration per flower head was monitored. Each beetle was observed for a total time of 30 min and 45 beetles were used per chemotype combination, with at least 10 responsive beetles per combination. Each beetle was tested only once.

### Weight gain assay

To evaluate the weight gain of beetles when feeding on pollen of the different chemotypes, we performed a no-choice assay. Pollen (0.145 g) pooled from several plants of one chemotype was inserted in a medium of agar (0.18 g, boiled in 10 mL water). The agar was filled in a Petri dish and stored at 4 °C until use. For the feeding assay, a disc (4 mm diameter) of the medium was put into a reaction tube with a small hole in the cap. Beetles were starved for 24 h, weighed (Sartorius microbalance, 0.001 mg; Germany) and then placed individually in the tubes. Tubes were placed in a climate chamber under a 16:8 light:dark cycle, 22 °C and 60% relative humidity. This experiment was replicated thrice, with each beetle tested once per chemotype, and the results were analyzed together in a mixed model. For each experiment, fresh medium was prepared. For each chemotype, 12-13 beetles were tested. The weight gain or loss of each beetle was expressed as a percentage change of weight (i.e., 0 % means no change in weight). After the assay, beetles were frozen and their sex was determined.

### Survival assay

To test the survival of the beetles on flowers of the three different chemotypes, beetles previously fed on tansy flower heads for a week were placed in white plastic boxes (22 cm x 13 cm x 12 cm), covered with a lid with gauze for ventilation. Inflorescences with freshly blooming flower heads were cut, provided in wetted foam in a falcon tube and placed in the boxes. The inflorescences were replaced every week, ensuring that pollen was available ad libitum. Boxes were cleaned regularly. Four boxes containing 15 beetles each were set up for each chemotype (total *n* = 60 per chemotype). Every three days, dead beetles were counted and removed. The observation time lasted until the last beetle had died.

### Statistical analyses

All analyses were performed using R v4.0.3 (https://www.r-project.org) and R Studio v1.4.1106 (https://posit.co/). Package stats (R Core Team, 2018) and ggplot2 (Wickham, 2016) were used throughout the analyses for several functions and graphs.

The terpenoid diversity of the flower heads and pollen of the chemotypes was analyzed using Renyi diversity profiles based on H_α_ = ln (Σ p_i_^α^)/1 - α, with α being the scale from 0 to infinity and p being the relative abundance of each terpenoid within an individual; applying the function ‘renyiresult’ of the package biodiversityR (Kindt, 2005). The Renyi profiles provide a range of diversity indices based on richness and proportional abundances (Tóthmérész, 1995). The scale parameter α is adjusted for different indices, with α = 0 for richness, α = 1 for Shannon diversity index, α = 2 for Simpson diversity index and α = inf for the Berger-Parker index (Tóthmérész, 1995). Higher α values place more weight on abundance over richness and evenness, with α = inf being related to the proportion of the most dominant species. When profiles do not intersect, the one with larger values of H_α_ is considered more diverse.

To test whether the composition of terpenoids, sugars and amino acids, each tested separately, differed among chemotypes, permutational multivariate analyses of variance (PERMANOVAs) were used (function adonis2 in package vegan; Oksanen et al., 2013). Data was Wisconsin-square root transformed to scale for the differences in compound variation. Chemotype and maternal origin of the plant were included in the model as fixed factors. The Kulczynski dissimilarity index and 1000 permutations were used in all PERMANOVAs. For post-hoc analyses of significant results, pairwise PERMANOVA comparisons were performed with a Benjamini-Hochberg (BH) correction of *p* values (package EcolUtils; Salazar, 2023). Visualisation of terpenoid profiles was done by projection onto a non-metric dimensional scaling (NMDS) plot (package vegan).

Total concentrations of sugars, total amino acids and total lipids were analyzed for differences across chemotypes. Data were first tested by using a Shapiro-Wilk test (package stats) for a normal distribution followed by Levene’s test for homoscedasticity of variances (package car; Fox, 2019). Data that met both conditions were subject to ANOVAs followed by Tukey’s post-hoc tests, if significant, while data that did not were tested using Kruskal-Wallis tests followed by Dunn’s post-hoc tests.

The preferences of the beetles in the olfaction and contact choice assays were analyzed using two-sided binomial tests (package stats). T-tests were used to compare the differences in the time spent in each arm in the contact choice assay. For the weight gain assay, a linear model was used to estimate the relation between the response variable, % weight gain of each beetle, and the predictor (fixed factor) variable chemotype (package lme4). Sex of the beetle and trial number were used as covariates and beetle number was used as a random effect. Likelihood ratio (chi-square) ANOVAs were used to derive *p*-values for the fixed effects and covariates. Post-hoc analyses on the factor chemotype were carried out using Tukey-corrected contrast tests on the least-squares means. Beetle survival on each chemotype was compared with an overall log-rank test, followed by pairwise log-rank tests with Benjamini-Hochberg correction for *p*-values and illustrated with Kaplan-Meier curves (package survminer; Kassambara, 2020).

## RESULTS

### Differences in terpenoid composition of flower heads and pollen among chemotypes

The major terpenoids of the flower heads mirrored those of the leaves, based on which the chemotypes had been defined, and showed a similar order of Shannon diversity indices (Supplement Table S1, Fig. S3). The Renyi diversity profiles revealed that the flower heads showed decreasing diversity from the Keto over the BThu to the Myrox chemotype, while this trend was reversed in the pollen (Fig. 1A, C). However, chemical richness values (represented by H_α_ = 0) were consistently higher in the BThu chemotype and lower in the Myrox chemotype.

**Fig. 1.**
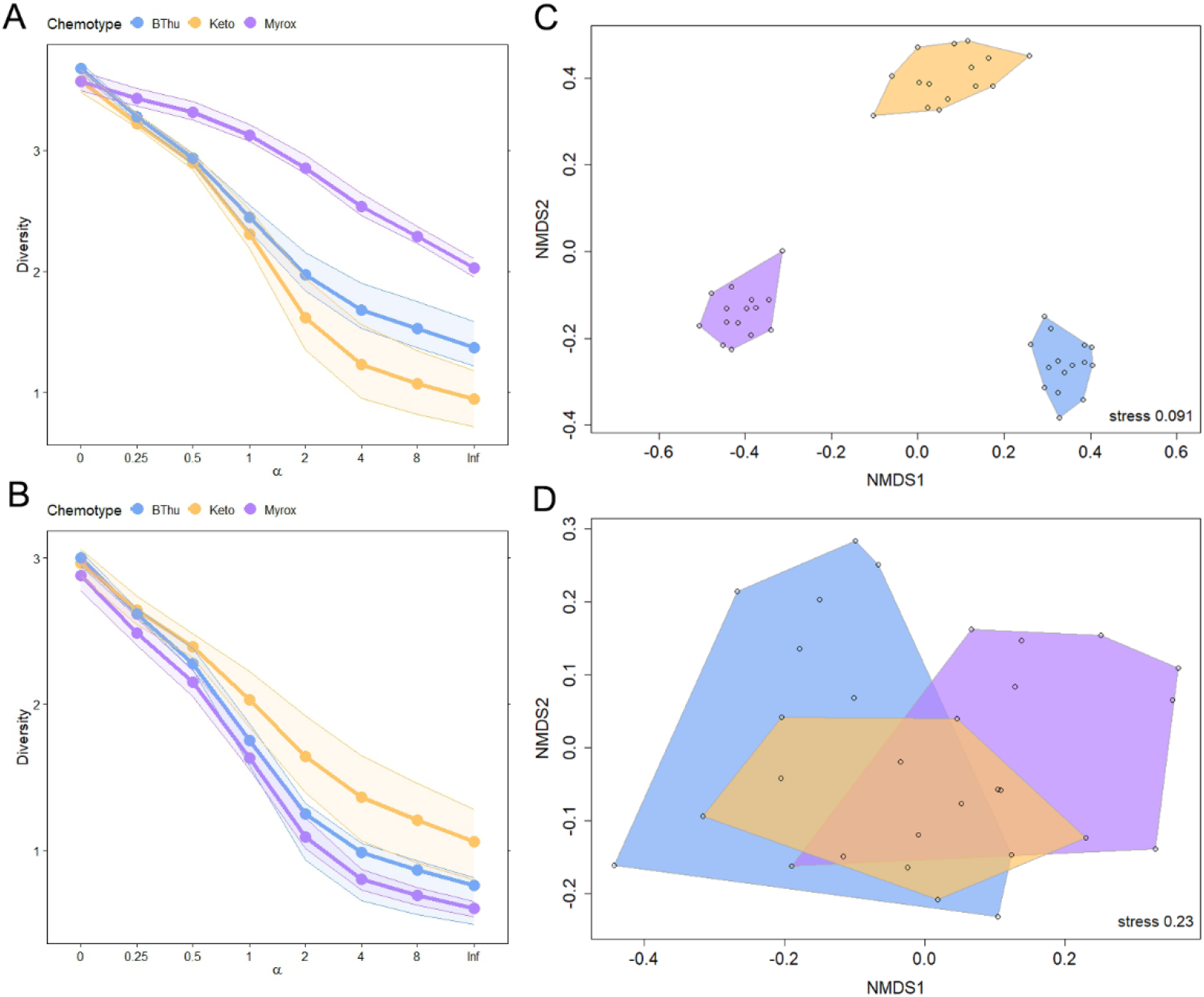
Terpenoid diversity and composition of flowers and pollen of *Tanacetum vulgare* plants belonging to the three chemotypes; yellow = Keto: artemisia ketone chemotype, blue = BThu: β-thujone chemotype, purple = Myrox: (*Z*)-myroxide-santolina triene-artemisyl acetate chemotype. Renyi diversity indices (H_α_) of flower heads (A, *n* = 15 per chemotype) and pollen (B, *n* = 8-13 per chemotype) with α = 0 is for species richness, α = 1 for Shannon index, α = 2 for inverse Simpson index (1/D) and α = Inf for Berger-Parker index. and NMDS plot of flower heads (C) and pollen (D) based on Kulczynski distances with *n*_permutation_ = 1000 and seed = 100.

The flower head terpenoid composition differed significantly among all three chemotypes (PERMANOVA, *F* = 37.93, *p* < 0.001; Fig. 1B) and also due to maternal origin (*F* = 1.413, *p* = 0.03). All three chemotypes differed from each other (*F_BThu-Keto_* = 29.38, *F_Keto-Myrox_* = 31.35, *F_BThu-Myrox_* = 41.15, *p_adj_* < 0.001 for all pairs). The terpenoid composition of the pollen also differed among the chemotypes (PERMANOVA, *F* = 2.73, *p* < 0.001; Fig. 1D). Pairwise comparisons revealed significant differences between the terpenoid profiles of pollen from the BThu and Keto (*F* = 2.43, *p* = 0.013) and the BThu and Myrox chemotype (*F* = 3.33, *p* = 0.003), while the pollen terpenoids of the Keto and Myrox chemotype were only marginally different (*F* = 1.97, *p* = 0.059). The total terpenoid content did not significantly differ between the chemotypes (Kruskal-Wallis test, *Χ^2^* = 5.003, *p* = 0.08; Fig. S4).

### Differences in nutrient composition of pollen among chemotypes

Five major sugars were found in the pollen, fructose, glucose, sucrose, an unidentified sugar alcohol (identified as such based on the *m*/*z* fragments 147, 217, 265, 305 and 318) and an unidentified trisaccharide (based on *m*/*z* fragments 147, 191, 204 and 217 as well as the retention time). The pollen sugar composition did not significantly differ among the chemotypes (PERMANOVA, *F* = 1.185, *p* = 0.34; Fig. 2A), but was significantly affected by the maternal origin (*F* = 2.106, *p* = 0.024). Fructose was the major sugar of the pollen. Likewise, none of the sugars and sugar alcohols did differ significantly among the chemotypes (ANOVA or Kruskal-Wallis test, *p* > 0.05). The unidentified trisaccharide was present only in trace amounts in the Keto and BThu chemotype (Fig. S5). The overall sugar content did not differ between the chemotypes either (Kruskal-Wallis test, *Χ^2^ = 0.638*, *p* > 0.05; Fig. S4).

**Fig. 2.**
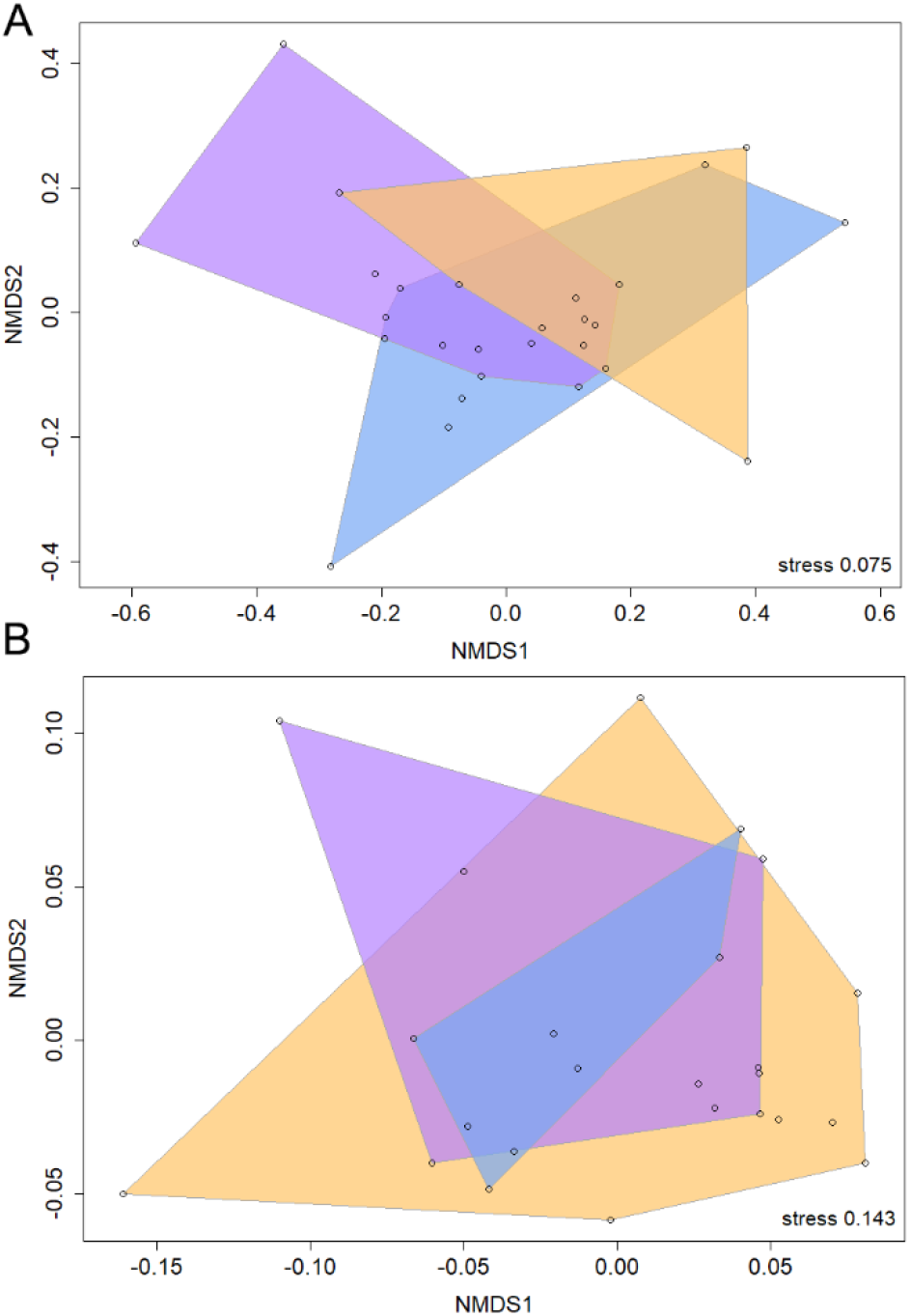
Sugar (A) and amino acid composition (B) of pollen of *Tanacetum vulgare*, displayed in NMDS plots based on Kulczynski distances matrices; yellow = Keto: artemisia ketone chemotype, blue = BThu: β-thujone chemotype, purple = Myrox: (*Z*)-myroxide-santolina triene-artemisyl acetate chemotype; *n* = 6-13 per chemotype for sugars and *n* = 6-10 per chemotype for amino acids. The NMDS analysis was carried out by scaling using Kulczynski distance matrices with *n*_permutation_ =1000 and seed = 100.

Nineteen amino acids were detected in the pollen, some of which were only present in trace amounts. The major amino acids included aspartate, proline, glycine, arginine, serine, glutamate and lysine (Fig. S6). The pollen amino acid composition did not significantly differ among the chemotypes (PERMANOVA, *F* = 0.393, *p* = 0.90; Fig. 2B) or due to maternal origin (*F* = 0.594, *p* = 0.94). There were also no significant differences in the amount of any of the individual amino acids among the chemotypes (ANOVA or Kruskal-Wallis test, *p* > 0.05). Likewise, the overall amino acid content did not differ among the chemotypes (Kruskal-Wallis test, *Χ^2^*= 0.461, *p* > 0.05; Fig. S4). The same was true for the total lipid content (ANOVA, *F* = 1.652, *p* = 0.21; Fig. S4).

### Olfactory and contact preferences of beetles towards chemotypes

Flower heads of the different chemotypes did not differ in attractiveness towards the beetles when compared to blank controls, except for the Keto chemotype, where beetles showed a marginal preference towards the blank control (binomial test *p* = 0.052, *n* = 60; Fig. 3A). In pairwise tests between chemotypes, the BThu chemotype was preferred over the Myrox chemotype (*p* = 0.047), while no preferences were found in the other combinations (Fig. 3B).

**Fig. 3.**
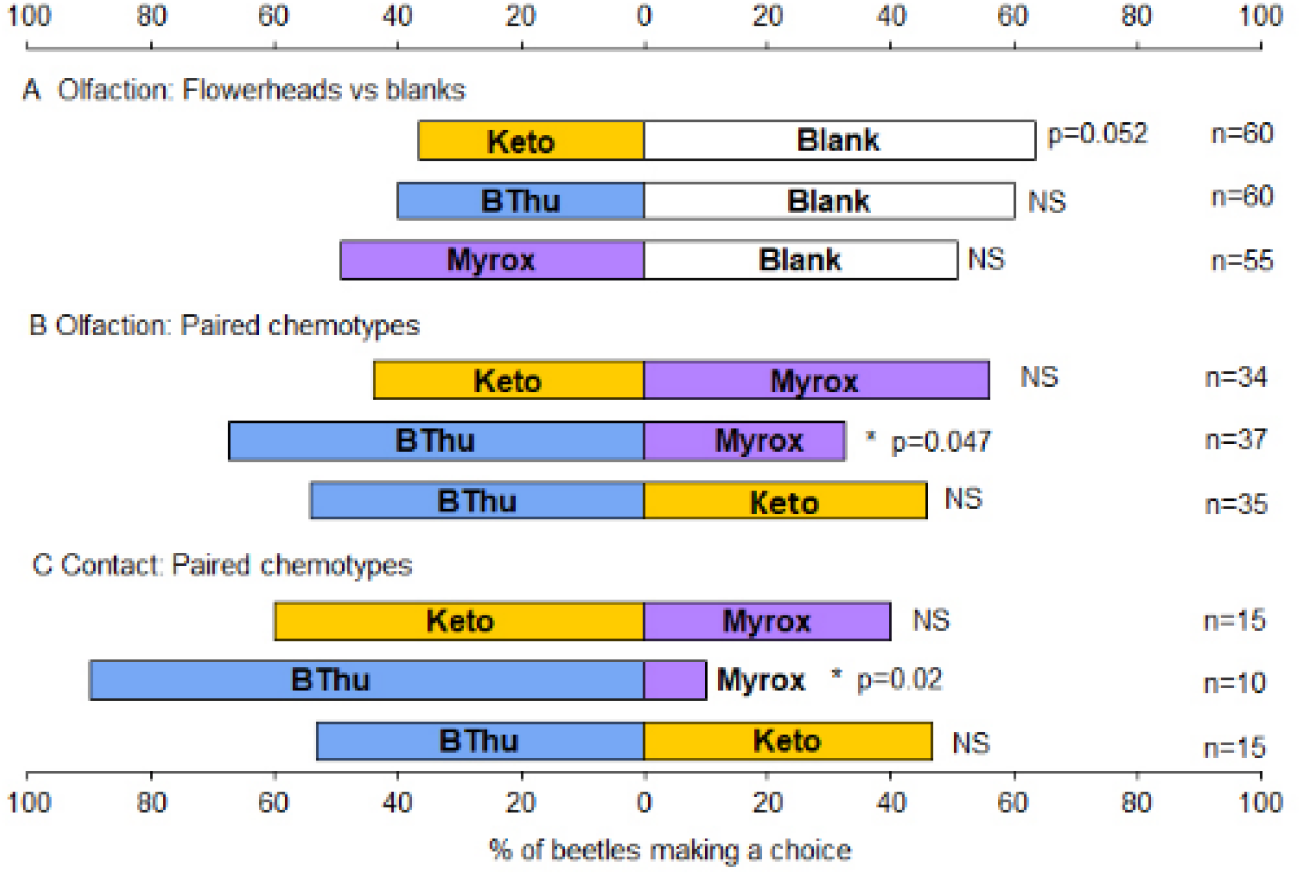
Choice behaviour of *Olibrus aeneus* beetles towards flower heads of different *Tanacetum vulgare* chemotypes, tested in olfactometer assays (A and B) and contact assays (C). Bars represent % of beetles tested for a pair of choices selecting a choice; Keto: artemisia ketone chemotype, BThu: β-thujone chemotype, Myrox: (*Z*)-myroxide-santolina triene-artemisyl acetate chemotype.

In pairwise contact choice assays, the BThu chemotype was significantly preferred over the Myrox chemotype (*p* = 0.02, *n* = 10), while no other significant differences were found between the other chemotype combinations (for *p* > 0.05) (Fig. 3C). Moreover, beetles also spent significantly more time on flower heads of the BThu chemotype compared to the Myrox chemotype (t-test, *t* = 3.067, *p* < 0.01) but showed no differences among the other pairs (Fig. S7).

### Weight gain of beetles on the chemotypes

The weight gain of the beetles differed significantly depending on the pollen source (linear mixed effects model with likelihood ratio tests, *Χ^2^* = 7.69, *p* = 0.021). Sex of the beetle had no effect on the weight gain. On pollen of the BThu and Keto chemotype, beetles on average gained weight, while they lost weight when feeding on pollen of the Myrox chemotype. These differences were significant for the BThu and Myrox chemotype (Tukey-corrected pairwise contrast test on least-squares means, *p* < 0.05; Fig. 4).

**Fig. 4.**
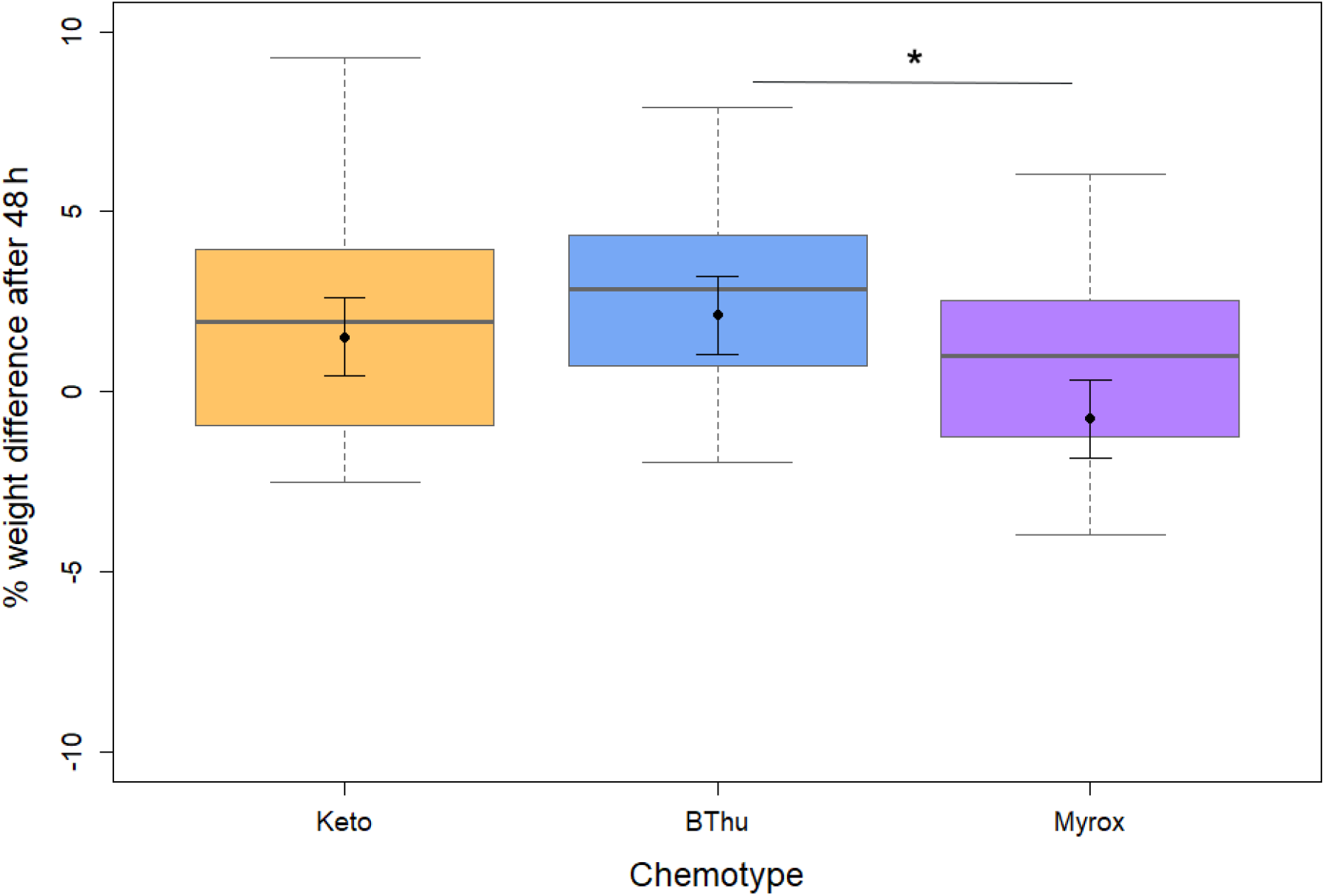
Weight gain of adult beetles of *Olibrus aeneus* (males and females pooled) after 48 h feeding on pollen of *Tanacetum vulgare* from different chemotypes; Keto: artemisia ketone chemotype, BThu: β-thujone chemotype, Myrox: (*Z*)-myroxide-santolina triene-artemisyl acetate chemotype. Box plots show median, lower (Q1) and upper (Q3) quartiles with whiskers, i.e., 1.5 times the inter-quartile range (Q3-Q1), *n* = 30-31 beetles per chemotype. LME model-derived means and standard errors are plotted over the box plots in black. Asterisk indicates significant difference (*p* < 0.05) in pairwise contrast tests.

### Survival of beetles on the chemotypes

In survival assays with adult beetles, mortality began as early as 3 days and showed high variation, with one beetle surviving up to 176 days from the start of the assay. Initial survival (within the first 25 d) dropped faster on the Myrox chemotype than on the other chemotypes. Across the entire period, the beetles showed different survival on the three chemotypes (log-rank test, *p* = 0.025; Fig. 5). Survival was highest on the BThu chemotype, and compared to BThu significantly lower on the Keto (pairwise log-rank test, *p* = 0.025) and marginally lower on the Myrox (*p* = 0.103) chemotype.

**Fig. 5.**
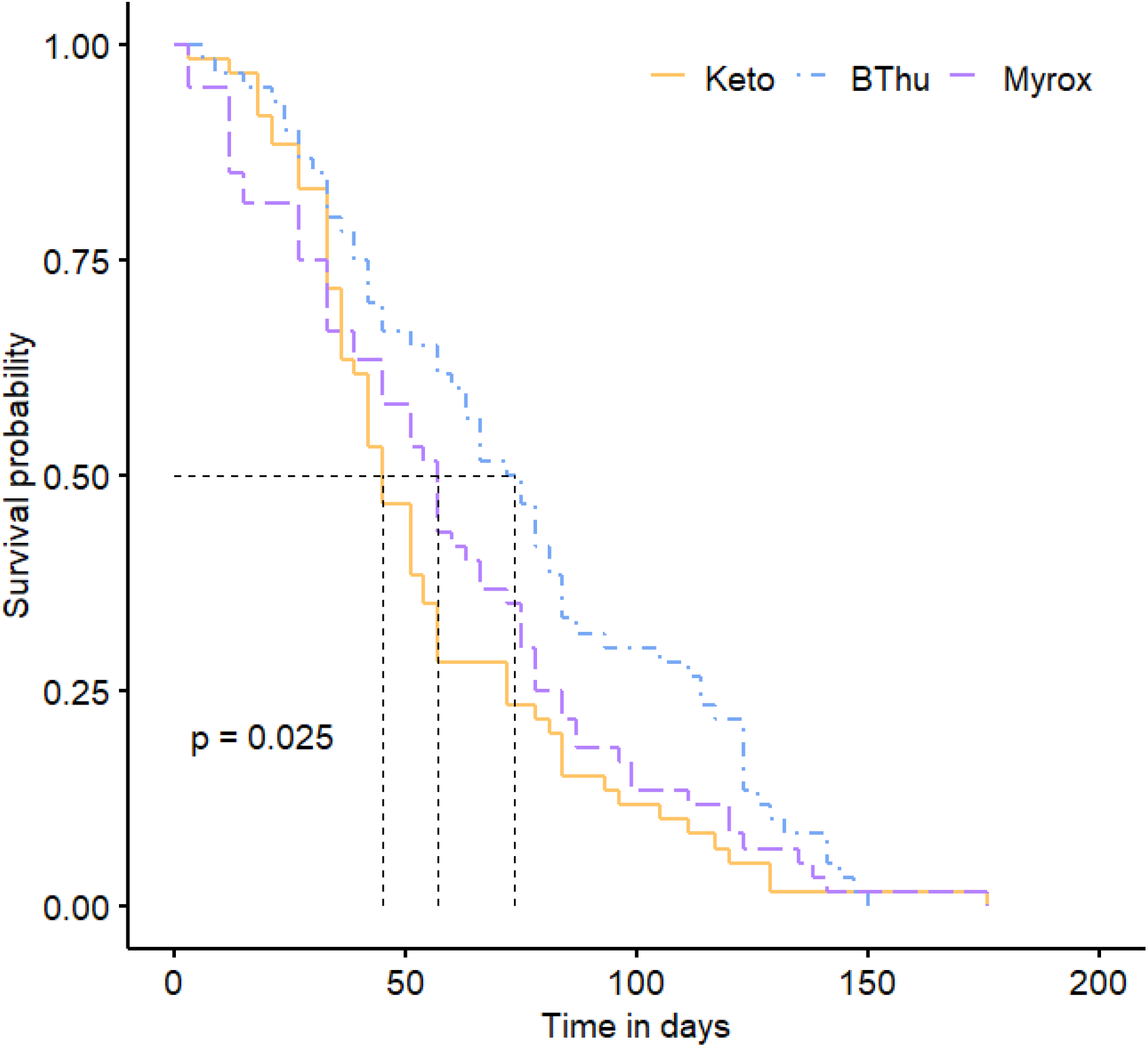
Kaplan-Meier survival curve of adults of *Olibrus aeneus* on flower heads of *Tanacetum vulgare* of the artemisia ketone chemotype, β-thujone chemotype and (*Z*)-myroxide-santolina triene-artemisyl acetate chemotype; initially *n* = 60 beetles per chemotype.

## DISCUSSION

Using various chemical analyses, our study revealed that within the highly chemodiverse plant *T. vulgare*, chemical differences among individuals were mostly found with regard to terpenoid composition of flower heads and pollen but less for pollen nutrients. Using bioassays, we demonstrated chemotype-specific consequences on the preference behaviour and performance of the oligophagous florivore *O. aeneus*.

The terpenoid composition of individuals of *T. vulgare* gives rise to unique chemotypes, which are mostly defined by the composition in the leaves (Clancy et al., 2016; Kleine and Müller, 2011; Ziaja and Müller, 2023), but occasionally also with regard to flower profiles (Judzentiene and Mockute, 2005; Eilers et al., 2021). As expected, the flower terpenoid profiles reflected the leaf chemotype, with the same predominant terpenoids characterizing the respective chemotype, as also found in previous studies (Judzentiene and Mockute, 2005; Eilers et al., 2021). Terpenoid synthesis is genetically determined (Jia et al., 2022), as are the resulting chemotypes of *T. vulgare* (Keskitalo et al., 2001; Lokki et al., 1973). However, interestingly the terpenoid profiles of flower heads and pollen differed in their order of diversity; flower heads of the Myrox chemotype had the highest and those of the Keto chemotype the lowest overall diversity, while the reverse order was found in the pollen. In contrast, the BThu chemotype showed the highest chemical terpenoid richness in both flower heads and pollen. Pollen is known to contain specialized metabolites, but the diversity and concentration compared to other floral and vegetative tissues is not well explored (Palmer-Young et al., 2019; Rivest and Forrest, 2020; Sasidharan et al., 2023). Terpenoids are probably not produced in the pollen but biosynthesized and stored within other floral tissues such as the anther glands (Goodger et al., 2021) or glandular trichomes (Ferreira and Janick, 1995). The terpenoids may then be released into or absorbed by the pollen during microsporogenesis, i.e., pollen formation.

The nutrient composition of sugars and amino acids was comparable in the pollen among the chemotypes, and may be more conserved due to selection by pollinators (Nicolson, 2011; Palmer-Young et al., 2019). Thus, our hypothesis that higher nutritional contents would be defended by more chemodiverse pollen could not be supported. While there were no differences in the amounts of nutrients in pollen at the chemotype level, an impact of maternal origin on pollen sugar levels was revealed. Likewise, in the leaf metabolite composition, differences due to maternal origin have been revealed (Dussarrat et al., 2023). These findings highlight that parental genotype can contribute to intraspecific chemodiversity.

The distinct chemical composition of the flower heads and pollen had a significant impact on the preferences and performance of adults of *O. aeneus*. Preferences represent the choice of an insect herbivore when provided with multiple food options. Volatile cues are usually the first line of floral cues important for flower visitors, potentially together with visual cues. Volatile and visual cues are particularly important for florivores, which have low mobility (Theis and Adler, 2012; Underwood et al., 2020). As a result, florivores may have also evolved host-tracking based on floral scents, as pollinators did (Schiestl, 2010; Schiestl, 2015; Strauss and Whittall, 2006), but with an unknown degree of specialization. In our experiments, the beetles did not show a strong response to flower head volatiles offered against blanks, but showed a significant preference for flower head volatiles of the BThu chemotype in choice assays against other chemotypes. The terpenoid richness was highest for this chemotype, indicating that terpenoid richness may be an important factor, but the beetles may also just have clear preferences for certain terpenoids of the BThu chemotype. The less attractive Myrox chemotype had an overall higher terpenoid diversity and (*Z*)-myroxide and santolina triene, the chemotype-defining compounds in the Myrox chemotype, are likely more volatile as they have higher estimated vapor pressures (Fig. S3) than several of the other detected terpenoids. These factors may have also influenced the olfactory responses of the beetles.

Contact cues, through gustation, are the second line of floral signals influencing preferences of florivores. Terpenes are often deterrent to flower visitors at the gustatory level (Theis and Lerdau, 2003). Moreover, chemical richness is known to have an influence on herbivory in plants, often leading to decreased herbivory (Theis and Lerdau, 2003). However, in our contact choice assays, BThu was found to be preferred over the Myrox chemotype even at the level of contact. Therefore, a high chemical richness may not necessarily deter floral visitors at the gustatory level. Flowers with high richness, along with low or intermediate Renyi diversity, as found in the BThu chemotype, might in fact be preferred by some flower visitors. Across different plant species, volatile richness may be linked overall to low toxicity in the pollen or to high nutrient content (Sasidharan et al., 2023). These traits could also be important towards attracting florivores through allowing detection of more olfactory cues from food such as pollen. Responding to more terpenoids may be particularly important for oligophagous florivores that could use richness to detect different host plant species.

In line with the preference for flower heads of the BThu chemotype, the performance assays revealed that *O. aeneus* performs well on this chemotype both in terms of weight gain within 48 h (in relation to the Myrox chemotype) and long-term survival (in relation to both the Keto and Myrox chemotype). Our hypotheses were thus partially supported, as feeding on the intermediate-diversity BThu chemotype resulted in the best performance. Chemical richness, highest in BThu, was not detrimental to feeding. Low overall diversity in the pollen of the Myrox chemotype might lead to greater toxicity at a given total terpene concentration (Eckhardt et al., 2014; Wetzel and Whitehead, 2020), if the fewer terpenoids are more abundant and thus, their individual concentrations are not “diluted”. Beetles feeding on the Keto chemotype with a high overall diversity showed also a poor long-term survival, potentially due to accumulating effects of terpenoids acting synergistically toxic over longer periods. Unfortunately, we did not observe oviposition and acknowledge that outcomes on larval performance or fecundity may be different. For example, larvae of honeybees were found to be more sensitive to specialized metabolites of pollen than the adults (Lucchetti et al., 2018).

Preference and performance do not necessarily go hand-in-hand (Charlery de la Masselière et al., 2017; Gripenberg et al., 2010). However, in our study, both were higher on the chemotype having higher chemical richness and intermediate overall diversity. These effects were consistent across the different assays. The beetles used in this study were collected mainly from *M. reticulata* individuals, which themselves show intraspecific variation in terpene diversity (Mežaka et al., 2020; Piri et al., 2019). Furthermore, during maintenance in the lab and prior to the bioassays, the beetles were exposed to all three tested *T. vulgare* chemotypes. In that way we avoided prior priming on a certain chemotype. In the field, beetles may also show a higher performance, i.e. higher weight gain and longer survival, when feeding on a mixed diet, as revealed in other insect herbivores (Bukovinszky et al., 2017; Friedrichs et al. 2022; Hagele and Rowell-Rahier, 1999). However, the impacts of chemodiversity on diet mixing are until now underexplored. In conclusion, our results highlight that both richness and overall diversity are important factors when determining chemodiversity of individual plants and their consequences on interacting insects.

## AUTHOR CONTRIBUTIONS

RS, EJE, and CM developed the research questions, study design and methods and EJE and CM acquired funding. RS and LB optimized and conducted the experiments. RS analyzed the data, prepared the figures and wrote the first version of the manuscript. All authors revised and commented on the manuscript.

## ACKNOWLEDGEMENTS

We thank Honey Seawon Yi for help with some bioassays, Manfred Krämer for assistance with beetle identification, Rabea Schweiger for help with chemical analyses and Pragya Singh and Silvia Eckert for help with data analyses. The olfactometer was constructed with help from Annette Beune (Glass Workshop, Chemistry Faculty) and the Mechanical Workshop of Bielefeld University.

