## Supplement for "Chemodiversity in flowers of *Tanacetum vulgare* has consequences on a florivorous beetle": Sasidharan_et_al_Supplement_20230619.pdf

**Supplementary Data**

**Table S1**: Shannon diversity ($H_s$) and richness of terpenoid profiles of *Tanacetum vulgare* leaves previously used for chemotyping and flower heads collected from greenhouse plants for the three chemotypes. Means for leaf samples and mean +/- SD for flower heads and pollen. Keto: artemisia ketone chemotype, BThu: β-thujone chemotype, Myrox: (*Z*)-myroxide-santolina triene-artemisyl acetate chemotype.

| **Chemotype** | **Leaf (chemotyped in 2020)** | | **Flower head (2021)** | | **Pollen (2021)** | |
|---|---|---|---|---|---|---|
| | $H_s$ | richness | $H_s$ | richness | $H_s$ | richness |
| Keto | 1.15 | 30.13 | 2.31 ± 0.36 | 35.87 ± 3.70 | 2.03 ± 0.21 | 19.75 ± 4.13 |
| BThu | 1.26 | 30.1 | 2.45 ± 0.12 | 39.67 ± 2.74 | 1.76 ± 0.35 | 20.2 ± 1.69 |
| Myrox | 2.16 | 28.5 | 3.13 ± 0.12 | 35.87 ± 4.10 | 1.63 ± 0.16 | 17.92 ± 2.43 |

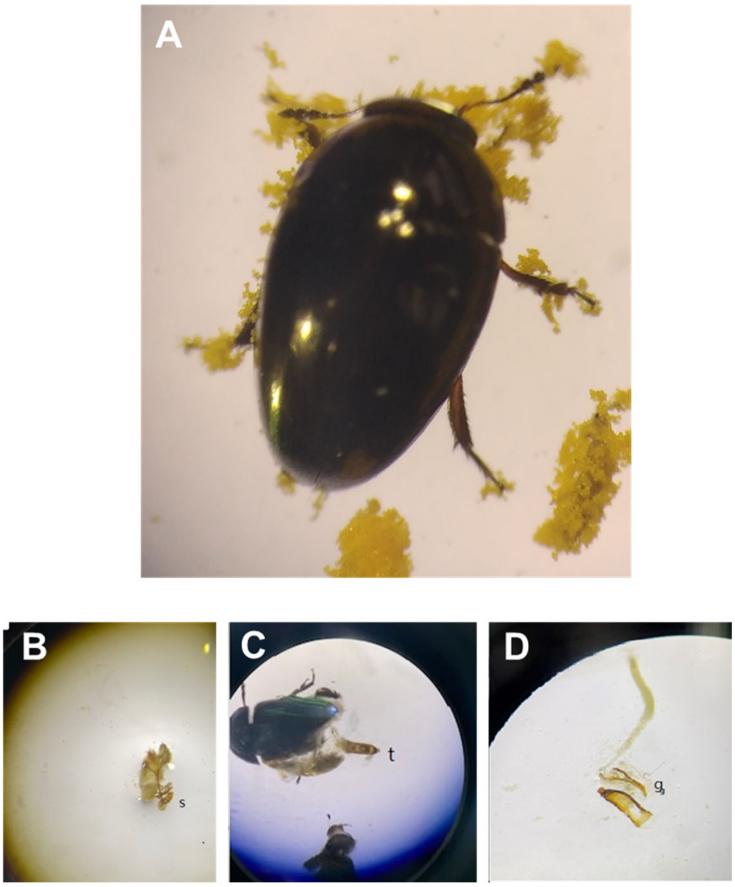


**Figure S1.** Images of (A) *Olibrus aeneus* showing characteristic features: olive-brown sheen, wide penultimate antennal segment, size 2.0-2.5 mm; (B-D) s= spermatheca, t= tegmen, g= female genitalia + ovipositor. The sex was determined based on the genitalia. The species was identified based on the Coleoptera database http://www.coleo-net.de.

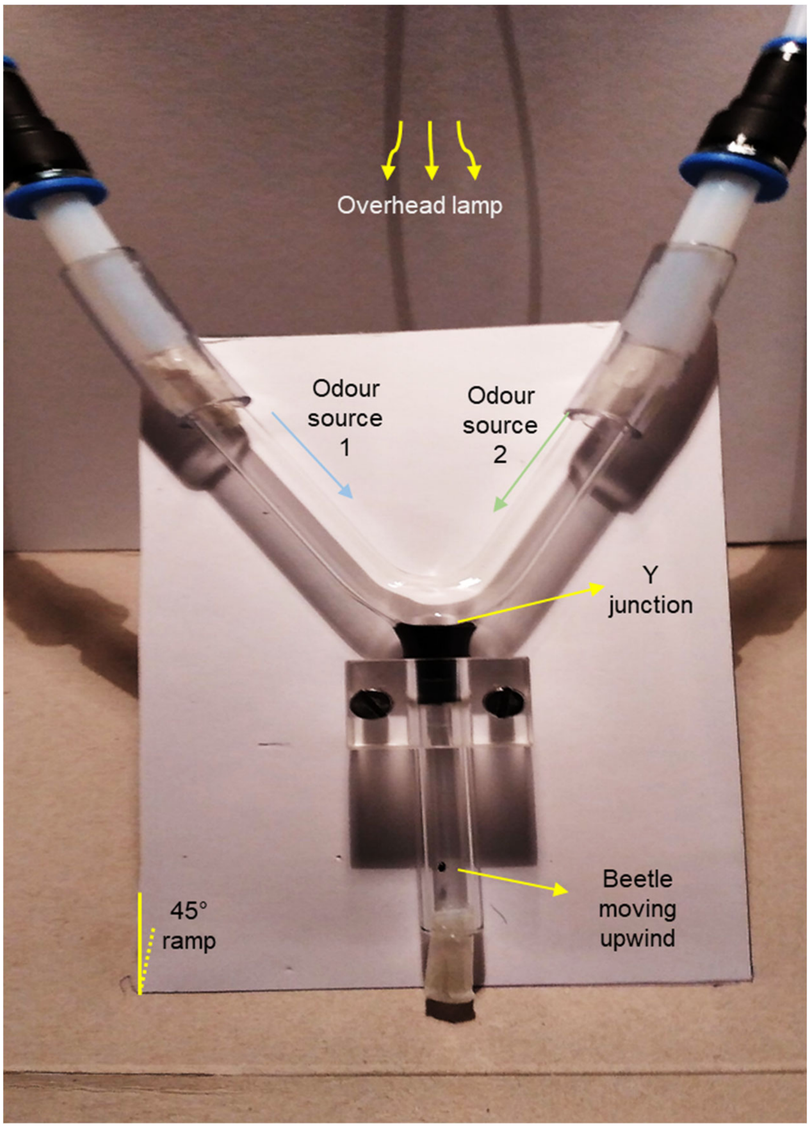


**Figure S2.** Y-tube olfactometer set-up used to test olfactory preferences of *Olibrus aeneus*. Details given in figure.

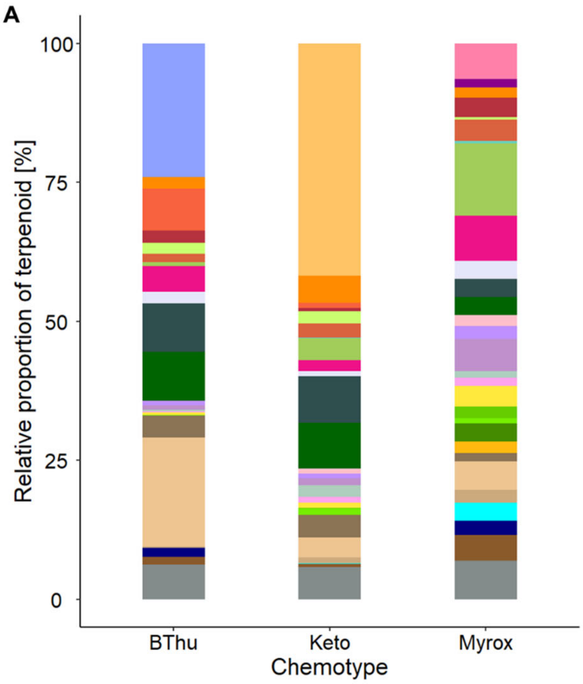


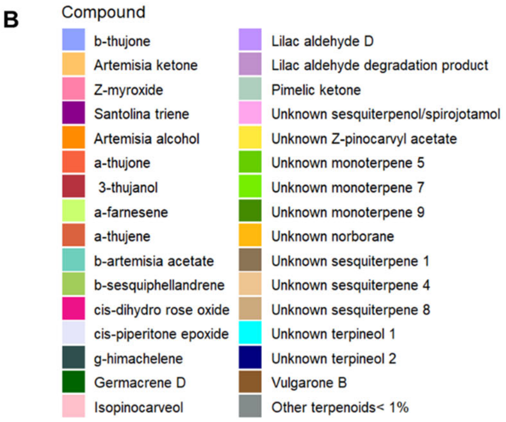


C

| **Terpenoid** | **Chemotype defined** | **Retention time in GC-MS (min)** | **Vapor pressure (mm Hg @ 25 °C)** |
|---|---|---|---|
| b-thujone | BThu | 12.26 | 0.3 (In-silico estimation, chemspider.com) |
| Artemisia ketone | Keto | 11.12 | 0.3 (In-silico estimation, chemspider.com) |
| Z-myroxide | Myrox | 9.83 | 1.39 (In-silico estimation, thegoodscentscompany.com) |
| Santolina triene | Myrox | 7.82 | 2.3 (In-silico estimation, chemspider.com) |
| Artemisia alcohol | Myrox | 11.52 | 0.035 (In-silico estimation, thegoodscentscompany.com) |

**Figure S3.** (A) Terpene profile of flower heads of *Tanacetum vulgare* of three chemotypes; BThu: β-thujone chemotype, Myrox: (*Z*)-myroxide-santolina triene-artemisyl acetate chemotype. (B) The terpenoids making up the profile are given in the key of which the (C) chemotype-defining terpenoids are provided in the table, along with their retention times and vapor pressures.

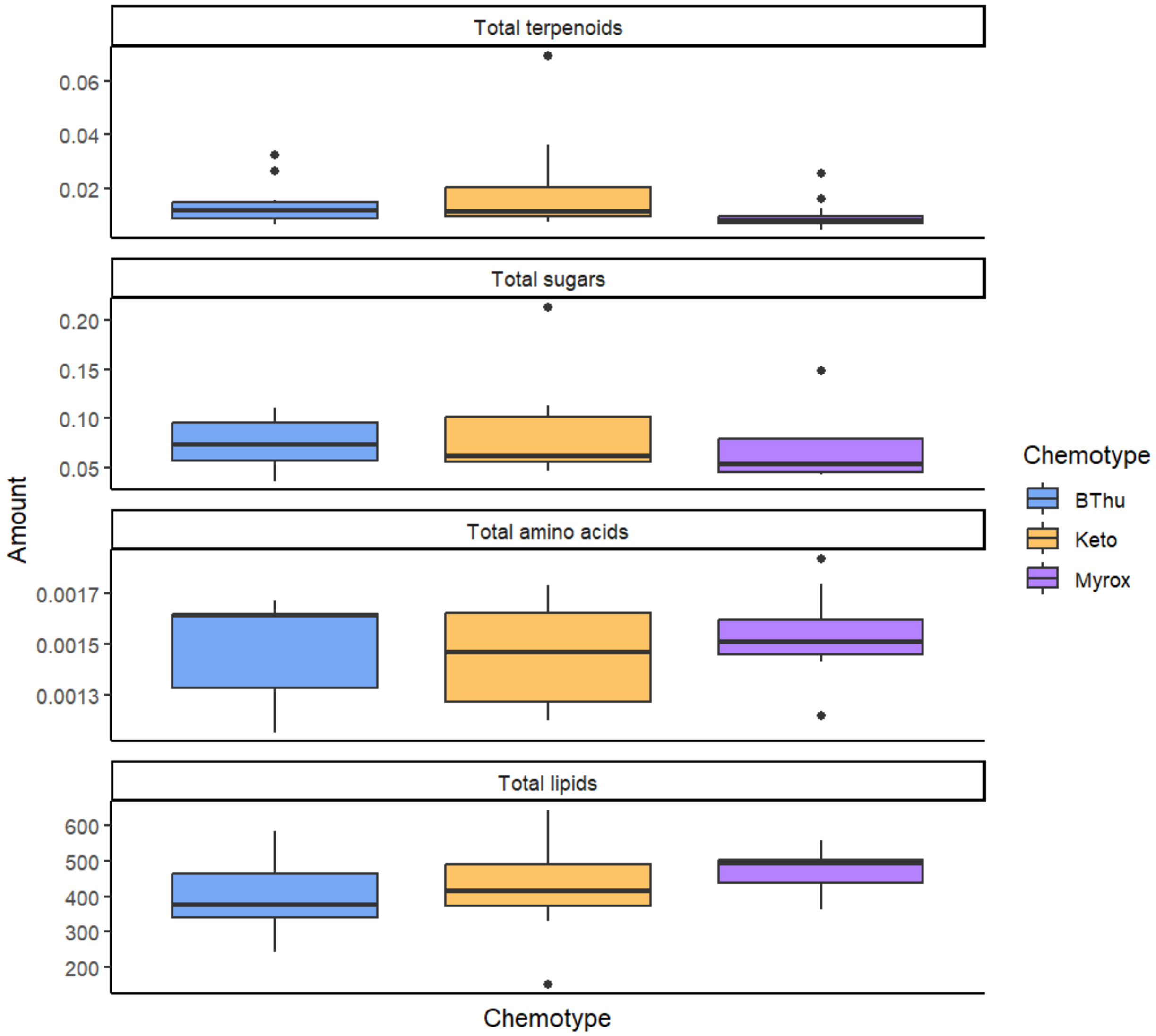


**Figure S4.** Total standardized amounts of terpenoids and nutrients in pollen samples of different chemotypes of *Tanacetum vulgare*; BThu: β-thujone chemotype, Keto: artemisia ketone chemotype, Myrox: (*Z*)-myroxide-santolina triene-artemisyl acetate chemotype. No significant differences were found among chemotypes.

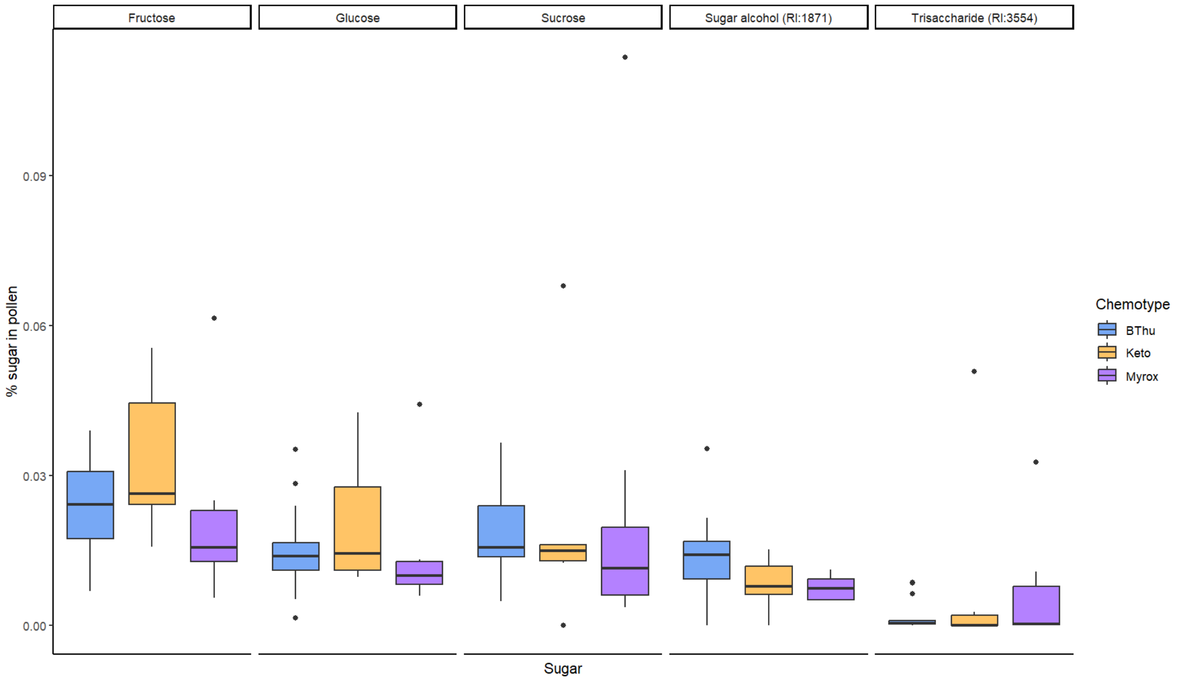


**Figure S5**. Total standardized amounts of sugars in pollen samples of different chemotypes of *Tanacetum vulgare*; BThu: β-thujone chemotype, Keto: artemisia ketone chemotype, Myrox: (*Z*)-myroxide-santolina triene-artemisyl acetate chemotype. No significant differences were found among chemotypes.

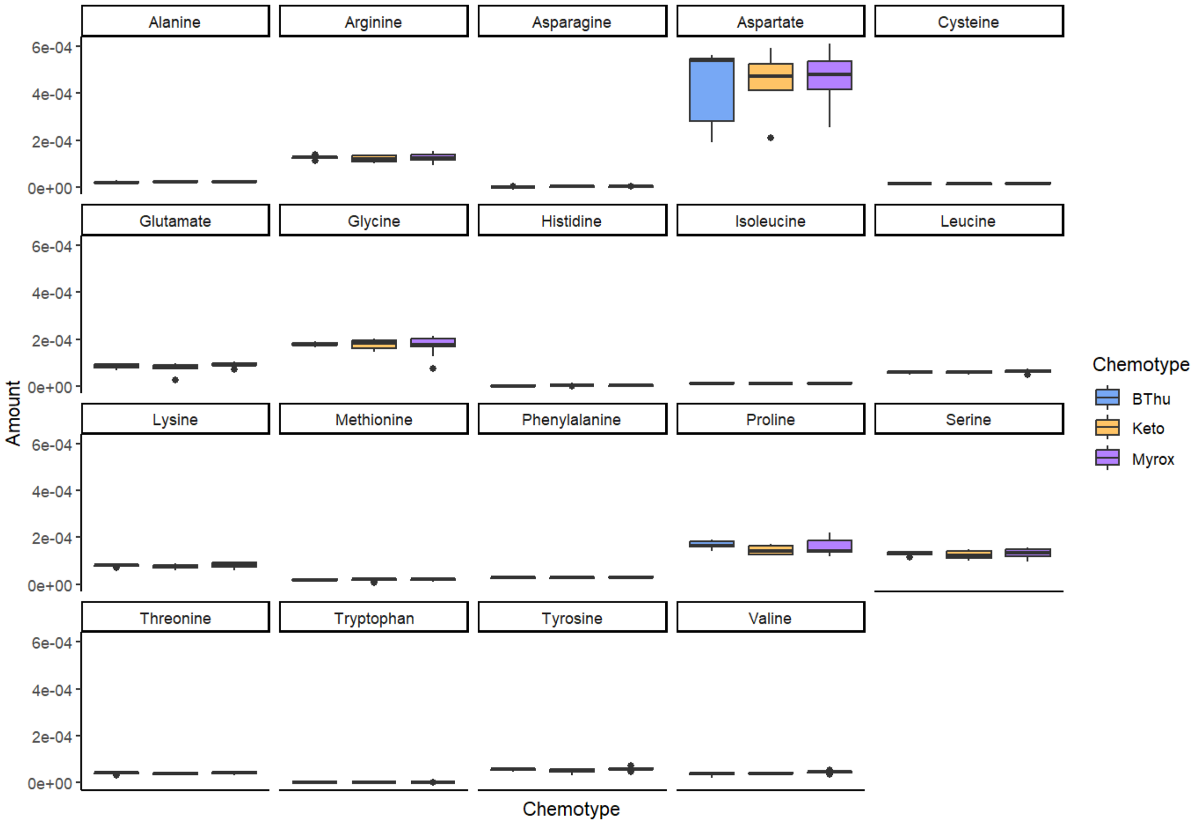


**Figure S6.** Total standardized amounts of amino acids in pollen samples of different chemotypes of *Tanacetum vulgare*; BThu: β-thujone chemotype, Keto: artemisia ketone chemotype, Myrox: (*Z*)-myroxide-santolina triene-artemisyl acetate chemotype. No significant differences were found among chemotypes.

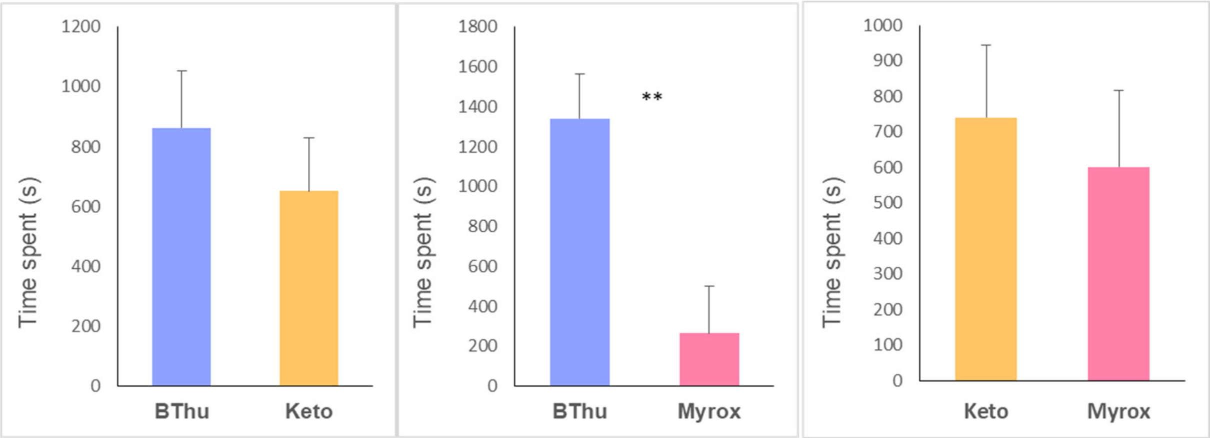


**Figure S7.** Mean (+SE) time spent by adults of *Olibrus aeneus* in each arm of a tube in a contact choice assay in which flower heads of *Tanacetum vulgare* of different chemotypes were offered; BThu: β-thujone chemotype, Keto: artemisia ketone chemotype, Myrox: (*Z*)-myroxide-santolina triene-artemisyl acetate chemotype. Pairwise comparisons, t-tests, ** indicates $p < 0.01$, $n$ = 10-15 replicates per beetle and chemotype combination.
